# Hypoxia induces extensive protein and proteolytic remodeling of the cell surface in pancreatic adenocarcinoma (PDAC) cell lines

**DOI:** 10.1101/2024.10.30.621099

**Authors:** Irene Lui, Katie Schaefer, Lisa L. Kirkemo, Jie Zhou, Rushika M. Perera, Kevin K. Leung, James A. Wells

**Author notes:** Indicates co-authors (Irene Lui, Katie Schaefer).

## Abstract

The tumor microenvironment (TME) plays a crucial role in cancer progression. Hypoxia is a hallmark of the TME and induces a cascade of molecular events that affects cellular processes involved in metabolism, metastasis, and proteolysis. In pancreatic ductal adenocarcinoma (PDAC), tumor tissues are extremely hypoxic. Here, we leveraged mass spectrometry technologies to examine hypoxia-induced alterations in the abundance and proteolytic modifications to cell surface and secreted proteins. Across four PDAC cell lines, we discovered extensive proteolytic remodeling of cell surface proteins involved in cellular adhesion and motility. Looking outward at the surrounding secreted space, we identified hypoxia-regulated secreted and proteolytically-shed proteins that are responsible for regulating the humoral immune and inflammatory response and an upregulation of proteins involved in metabolic processing and tissue development. Combining cell surface N-terminomics and secretomics to evaluate the cellular response to hypoxia enabled us to identify significantly altered candidate proteins which may serve as potential biomarkers and therapeutic targets in PDAC. Furthermore, this approach provides a blue print for studying dysregulated extracellular proteolysis in other cancers and inflammatory diseases.

## 1. Introduction

Pancreatic ductal adenocarcinoma (PDAC) represents more than 90% of pancreatic cancers. Over 50% of patients are diagnosed in the advanced stages of the disease when the tumor cells have metastasized^1^. A lack of detectable biomarkers and non-specific symptoms results in a delay in diagnosis and treatment. While treatments for many cancers have improved over the past few years, the 5-year survival rate for PDAC remains disappointingly low at 12.5%^1^. Understanding the cellular physiology of PDAC and its environment can lead to improved efforts to treat the disease.

The tumor microenvironment (TME) has a major impact on cancer progression, including effects on proliferation, invasion, metabolism, angiogenesis, metastasis, and immunosuppression^2,3^. Hypoxia is a widespread TME characteristic, caused by alterations in vascularization, which impacts cellular proliferation, metabolism and resistance to therapy^4–6^. Furthermore, hypoxic conditions create a hostile environment for immune cells and suppresses immunity^7–9^. Most tumor tissues range from 2-10 times more hypoxic than their respective normal tissues^4^, and strikingly, PDAC tumors can be as much as 25 times more hypoxic^4,10^.

Another hallmark of TME is the dysregulation of proteolysis^11^. Proteolysis is an irreversible, post-translational modification that contributes to many cellular and physiological processes, including cell signaling, protein processing, tissue remodeling, cellular migration, and programmed cell death to name a few^12–14^. Aberrantly-regulated, extracellular proteases contribute to the progression of disease by affecting cellular growth, responses to apoptosis and senescence, angiogenesis, invasion, metastasis, and inflammation^15,16^. To identify how proteases contribute to the remodeling of the cell surface and the surrounding extracellular milieu, several mass spectrometry (MS) methods have been developed to identify precise cleavage sites^17–22^ and shed or secreted proteins (secretome)^23–26^.

Previous studies have investigated the role of hypoxia in PDAC and its effect on various cellular phenotypes. For example, hypoxia is known to promote cell migration via *miR-150* downregulation^27^, and can contribute to the epithelial to mesenchymal transition (EMT)^28,29^. RNA analysis of patient samples^30,31^ and MS analysis of extracellular protein expression in mouse xenografts^32^ have provided insight on the proteomic landscape of PDAC tumors. However, it is poorly understood how a hypoxic environment changes cell surface proteolysis and the shedding and secretion of extracellular proteins in PDAC tumors.

To address how a hypoxic environment affects the proteolytic status of PDAC cell surfaces, we utilized a glycan-tethered stabiligase (GT-stabiligase) to identify cell surface proteolytic neo-N-termini across 4 PDAC cell types (KP4, Panc-1, PaTu8902, and MIAPaCa-2). Using secretomic methods ^23^, we also studied how hypoxia affects the shedding and secretion of proteins into the extracellular space. In total, we identified 906 unique N-termini that mapped to 384 cell surface proteins, and 517 secreted proteins. We found that hypoxia induces extensive remodeling of the proteolytic and extracellular protein landscape of PDAC, especially affecting proteins involved in cellular adhesion, motility, and response to the innate immune system. We believe this work provides important new targets which could serve as biomarkers or targets for therapies.

## 2. Results

### 2.1 Identifying cell surface neo-N-termini displayed on PDAC cells under hypoxic conditions

To globally identify precise cleavage sites across PDAC cells, we leveraged a cell surface proteolysis (CSP) technology that we previously developed^22^. In this approach, we employ stabiligase, an engineered ligase that preferentially labels N-terminal amines with a biotinylated-peptide ester substrate. Using glycan chemistry, we chemically-tether an aminooxy-conjugated stabiligase to native glycans on living cells, which improves the efficiency by >20-fold for selectively labeling of cell surface neo-N-termini for proteomic identification.

We selected four PDAC cell lines (KP4, Panc-1, PaTu8902, and MIA PaCa-2) which we cultured by stable isotopic labeling of amino acid (SILAC) media (**Figure 1a**) and then expanded under hypoxic (1% O_2_) or normoxic (5% O_2_) conditions for 72 hours. We then harvested cells, tethered GT-stabiligase to their cell surface glycans, and briefly pulsed cells with a peptide ester substrate featuring a biotin handle, a TEV-cleavage site, and an amino-butyric acid (Abu) mass tag. Labeled cells surface proteins were subsequently isolated and enriched on streptavidin, and digested on-beads with trypsin. Following a TEV protease treatment, an enriched pool of N-terminal peptides labeled with the Abu mass tag were then released from resin for proteomic identification using tandem mass spectrometry. Across the four cell lines, we identified 906 unique N-terminal peptides that mapped to 384 membrane proteins (**Figure 1b)**. Not surprisingly, the majority of identified proteins corresponded to type I single-pass transmembrane proteins, that reflects their relative abundance on the cell surface (58%) and their extracellular N-terminal outside orientation. The remaining N-termini were distributed across multi-pass proteins (22%), secreted proteins (10%), glycophosphatidylinositol (GPI)-anchored proteins (7.4%), and finally, type II transmembrane proteins with a C-terminal outside orientation (1.6%). We found that the vast majority of N-termini (88%) mapped to the extracellular domains corresponding to extracellular proteolysis sites (neo-N-termini); the remaining peptides are associated primarily with protein trafficking and maturation such as signal peptide cleavage sites, initiator methionine removal, and pro-domain removal. Using PDB deposited structures (**Figure 1d**) and AlphaFold 2.0 predicted structures (**Figure S1**), the secondary structure surrounding extracellular cleavage sites correlated mostly with loops/unstructured domains, consistent with the substrate preferences of proteases. Additionally, we mapped the amino acid distances between the neo-N-termini and the membrane anchor of the corresponding protein (**Figure 1e**), which revealed a distance distribution indicating that while many cleavages correlate with the shedding of whole protein ectodomains (within 40 amino acids of the membrane), a significant portion occurred away from the membrane.

**Figure 1:**
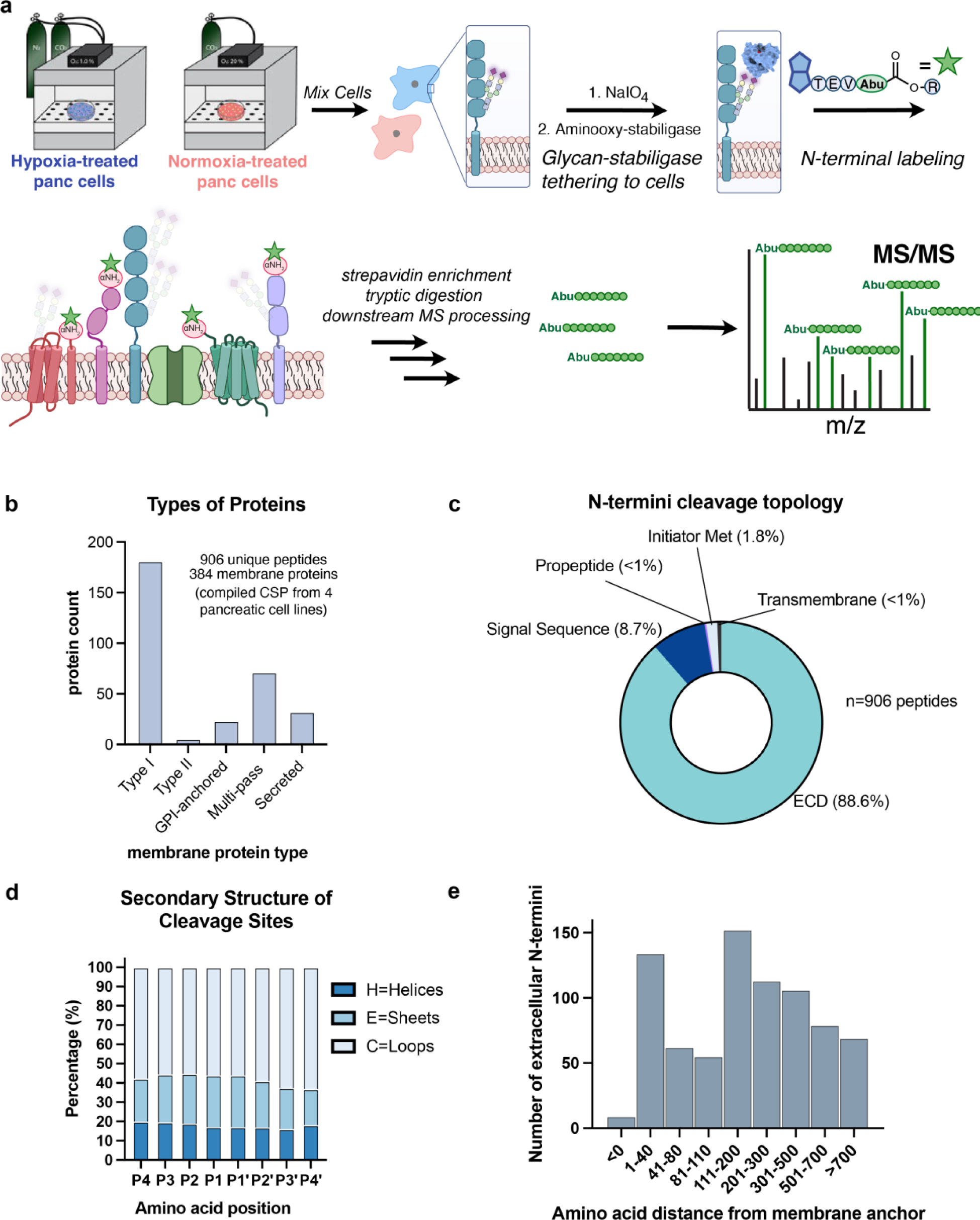
Glycan-stabiligase labels cell surface N-termini on PDAC cell lines under normoxic and hypoxic conditions. **a.** Schematic of procedure to identify N-terminal peptides using cell surface N-terminomics. PDAC cells are grown in heavy or light SILAC media under normoxic or hypoxic conditions. After mixing cells from each condition, GT-stabiligase was tethered to the cell surface. Tethered cells were then briefly incubated with a peptide substrate (Biotin-TEV cleavage site-Abu-ester-R) which was subsequently transferred to N-termini as catalyzed by stabiligase. For proteomic analysis, biotinylated proteins were enriched by streptavidin (SA), proteolytically digested, and N-terminal peptides were released upon TEV cleavage which results in aminobutyric acid (Abu) tagged peptides for LC-MS/MS identification. **b.** Across four cell lines we identified 906 unique N-termini corresponding to 384 membrane proteins. Of those annotated in UniProt, the majority of proteins identified were Type I single-pass transmembrane proteins, followed by a number of multi-pass and secreted proteins. **c.** Neo N-termini cleavage topology was analyzed and over 88% of N-termini mapped to extracellular protein loops. Cleavage events were also observed at regions that corresponded to initiator methionine removal, propeptide cleavage, signal peptide processing, and some cleavages within transmembrane regions. **d.** Using PDB deposited structures, predicted secondary structures for cleavage sites were made. H represents helix, E represents sheets, and C represents loops, with the majority of cleavages occurring in loops or unstructured areas. **e.** Distances between the cleavage site and the cell membrane were approximated based on the number of amino acids between the cleavage site and the membrane anchor.

### 2.2 Hypoxia causes widespread changes in proteolytic activity

Across the four PDAC cell lines, CSP data showed that there was not a general consensus proteolysis signature (**Figure S2a,** n=5), consistent with the known observation that common cancer phenotypes can arise from different molecular events. While each cell line demonstrated a unique proteolytic profile in response to hypoxia, Gene ontology (GO) analysis of biological processes of all peptides (**Figure S2b**) revealed an over-representation of proteins involved in receptor activity such as receptor tyrosine kinase (RTK) signaling, transducer and transmembrane signaling receptor activity. An over-representation of proteins involved in cell adhesion, migration, and locomotion were found based on GO analysis (**Figure 2c**). Hypoxia is known to affect cellular adhesion and migration^33–36^ and our data suggests that proteolytic remodeling of these molecules correlate with these phenotypic alterations.

**Figure 2:**
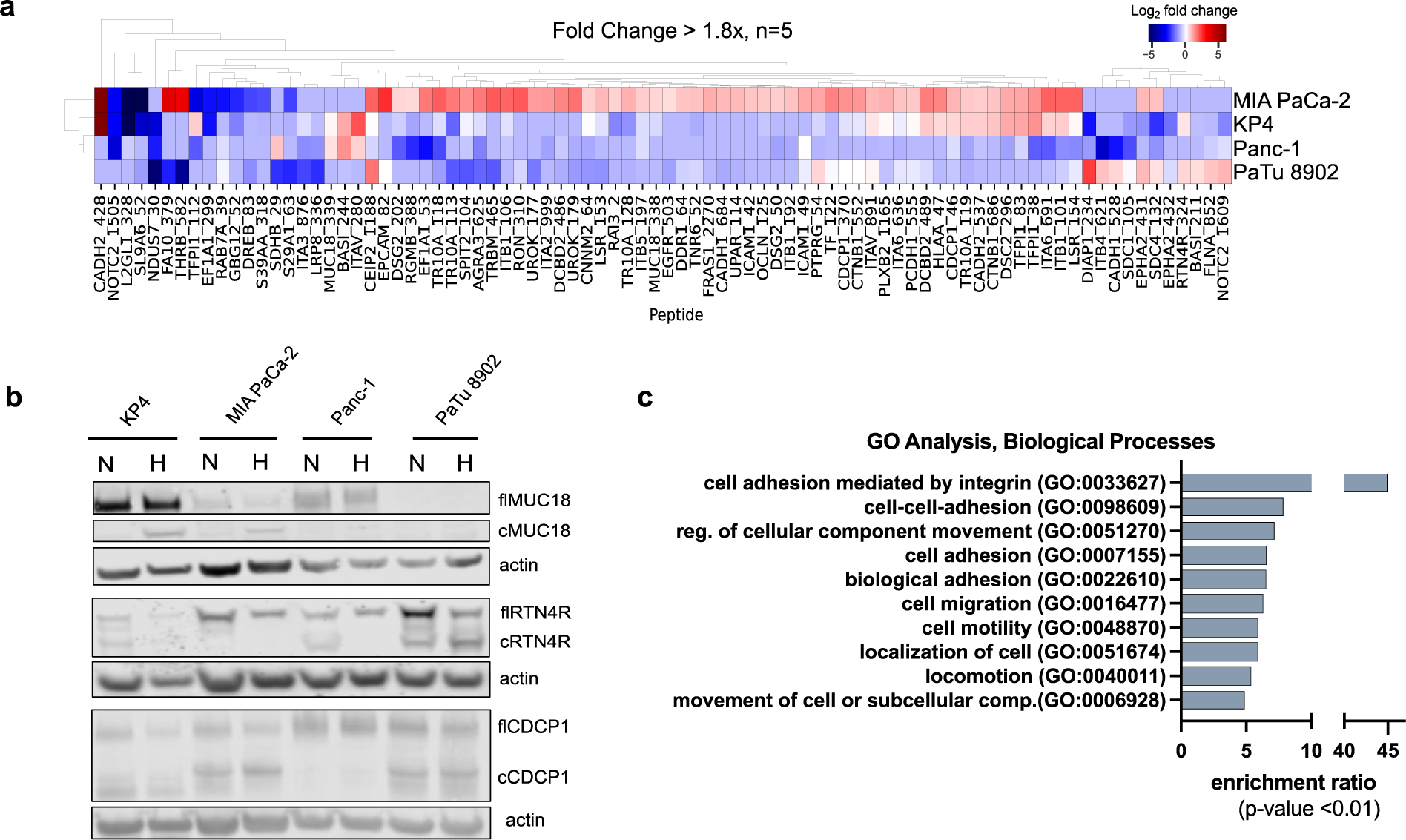
Hypoxic conditions induce extensive proteolytic remodeling on the cell surface. **a.** Heatmap of proteolytic neo-N-termini of membrane proteins showing significant up or down regulation (≥ 1.8-fold change) under hypoxic conditions compared to normoxic conditions. Peptides overrepresented under hypoxic conditions are represented in red while those with a higher presence in normoxic conditions are represented in blue. In KP4, Panc-1, and PaTu 8902 cells, overall surface proteolysis is decreased while MIA PaCa-2 cells showed a general increase. **b.** Western blot analysis of full length (fl) and cleaved (cl) isoforms of MUC18, RTN4R, and CDCP1 were consistent with quantitated proteolytic events identified by mass spectrometry. **c.** Gene ontology (GO) analysis of biological processes revealed that cleavages affecting proteins involved in cellular adhesion and migration were overrepresented under hypoxic conditions.

Across PDAC cells, we identified 84 neo-N-termini that had a greater than or equal to 1.8-fold change in abundance under hypoxic conditions in at least 2 cell lines (**Figure 2a**). Among these neo-N-termini, many peptides were less abundant under hypoxic conditions in the KP4, Panc-1, and PaTu 8902 lines. In contrast, MIA PaCa-2 cells demonstrated an increase in proteolysis at the same cut sites. We selected a panel of proteolyzed proteins for biochemical validation using western blot analysis. By analyzing the full-length (fl) and cleaved (c) proteoforms of membrane proteins MUC18, RTN4R, and CDCP1, we observed bands of proteoforms consistent with mass spectrometry analysis (**Figure 2b**). Notably, we observed several proteins with N-termini that differed only by a few amino acids. For example, the cell adhesion protein MUC18 cleavage is increased under hypoxia at residue 338 in MIA PaCa-2 cells, while showing slightly lower levels in the other cell lines. However, in KP4 and Panc-1 cells, a cleavage at residue 339 becomes more prominent and results in a similarly sized product identified by western blot. This likely reflects the degeneracy of some proteolytic sites, such as those observed in the cell surface receptor CDCP1 that is also abundantly shed on pancreatic cancer cells^59^.

### 2.3 Hypoxia induces changes in the secreted proteome

In addition to understanding how hypoxic conditions may alter the PDAC cell surfaceome, we also wondered how hypoxia may alter the secretome, the network of secreted and proteolytically-shed proteins from the cell surface. In CSP, we predominantly capture cell surface tethered proteins and we reason that the residence time of extracellular, soluble proteins near the cell surface may not be long enough to enable GT-stabiligase to label most secreted proteins. To evaluate secretome changes, we turned to a secretomics technology developed by the Lichtenthaler group^23^, outlined in **Figure 3a**. In this approach, hypoxic- and normoxic-cultured PDAC cells were metabolically labeled with N-azido-mannosamine (ManNaz) to incorporate azido-bearing sugars into secreted proteins. We then enriched secreted proteins away from the serum for mass spectrometry identification. Similar to the N-terminomics analysis, a majority of the 517 proteins identified in the supernatant were derived from type I single-pass transmembrane proteins (**Figure 3b**) as expected based on type I abundance. In fact, we found a larger number of annotated secreted proteins relative to cleaved proteins left on the membrane by N-terminomics suggesting the remaining fragment could be turned over more rapidly.

**Figure 3:**
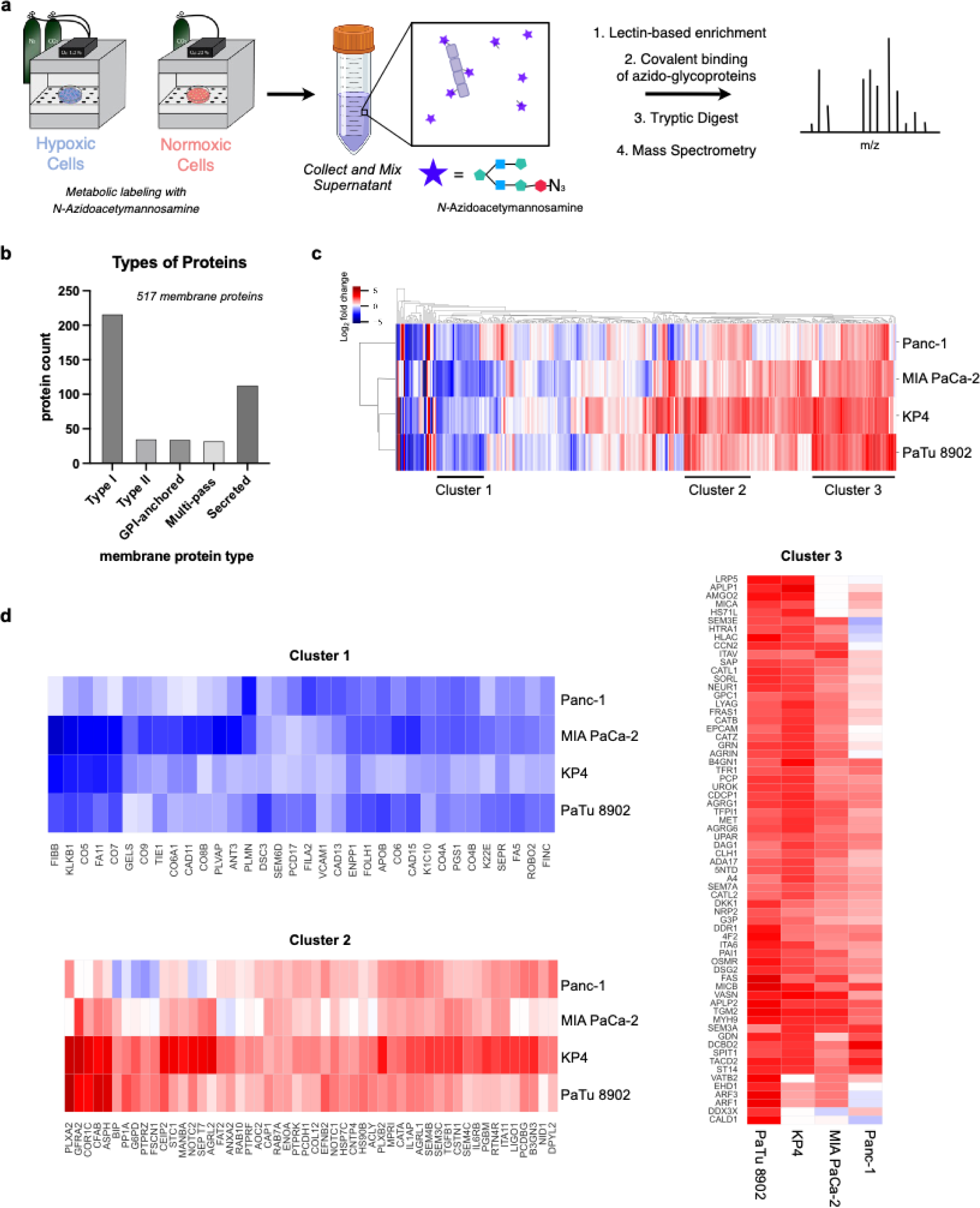
Analysis of the secretome of PDAC cell lines. **a.** Simplified schematic of the method to analyze secreted proteins. Cells are grown in heavy or light SILAC media under normoxic or hypoxic conditions and incubated with N-azido-acetylmannosame for 72 hours, resulting in azido-labelled glycoproteins. The supernatant from each condition was collected and mixed before lectin-based enrichment isolated all glycoproteins from the supernatant. Click chemistry was then employed to further enrich for azido-glycoproteins. Samples were digested with trypsin and peptides analyzed by mass spectrometry. **b.** 517 membrane proteins were identified; of those annotated, type I single-pass transmembrane proteins made up the largest portion of identified proteins, followed by secreted proteins, with a similar number of type II, multi-pass, and GPI-anchored identified. **c.** Global heat map analysis across four cell lines revealed three clusters of proteins. **d.** Cluster 1 contained proteins with decreased identification under hypoxic conditions in all four cell lines. GO analysis of biological processes (**Figure S3**) revealed a decrease in shed proteins involved in inflammatory and immune responses. Cluster 2 identified an increase in proteins involved in regulation of cellular metabolism. Cluster 3 showed an upregulation in proteins in tissue development and antigen processing and presentation.

A global examination of secreted proteins (**Figure 3c**) reveals three clusters of proteins across the four cell lines: Cluster 1 (**Figures 3d and S3a**) represents a group of downregulated proteins in hypoxia that, based on GO analysis, is linked to inflammatory and immune responses; Cluster 2 (**Figures 3d and S3b**) includes proteins that are slightly upregulated in hypoxia and are over-represented in molecules that regulate cellular metabolism; and Cluster 3 (**Figures 3d and S3c**) showed an increased number of shed/secreted molecules involved in antigen processing and presentation as well as those involved in endothelium and tissue development. Collectively, these data corroborate existing data that hypoxic conditions can aid in immune evasion^37,38^ and affect cellular metabolism to promote growth of the cancer cells^39–41^.

### 2.4 Proteolytic signatures are a consequence of many proteases

It is striking that we identified 36 extracellular proteases within the secretome samples (**Figure 4a**). We found 15 serine proteases, followed by ten metalloproteases and seven cysteine proteases plus a few others (**Figure 4b**). We observed upregulation of extracellular proteases such as ADAMs, cathepsins, and matrix metalloproteinases (MMPs) that are well-established to be involved in extracellular matrix degradation to increase tissue invasion. We also identified decreased abundance of proteases that are involved in coagulation and angiogenesis (*ex*., KLK2, CTSB, MMP2, **Figure 4a**).

**Figure 4:**
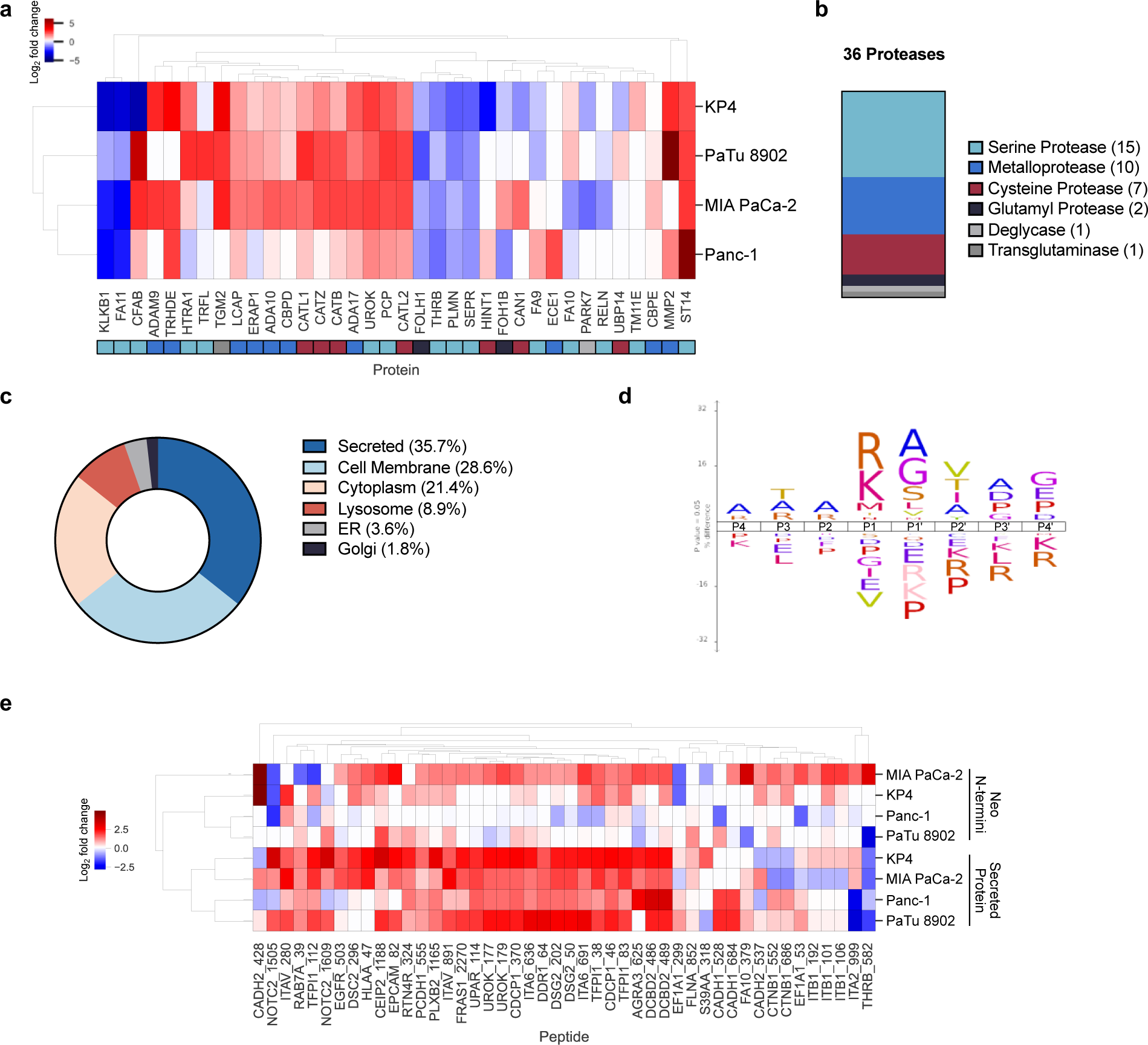
Secreted proteases and proteins under hypoxic conditions and correlation of N-termini and secretomics analysis. **a.** Proteases identified in cellular supernatant under hypoxic conditions across four cell lines. Colored tiles beneath the heat map annotate the type of protease, correlating with those in (**b**). **b.** Classification of identified proteases. **c.** Annotated location(s) for identified proteases according to Uniprot. **d.** iceLogo visualization of P4-P4’ residues flanking the cleavage site across four cell lines. **e.** Heatmap comparison between shared N-termini and corresponding secreted protein enrichments.

We analyzed the P4-P4’ amino acid motif IceLogo for the hypoxia-induced neo-N-termini (**Figure 4d**). The IceLogo did not indicate a definitive consensus sequence surrounding cleavage sites, which is not unexpected given the breadth of proteases identified here. Nonetheless, we did observe an overrepresentation of arginine and lysine in the P1 position that reflects the dominant number of serine proteases we identified that are known to prefer basic P1 residues.

Across both CSP and secretomics data, we identified 11 proteases common to both data sets (**Figure S4a**). In CSP, we identified neo-N-termini that corresponded to pro-sequence cleavage and proteolytic activation sites within the cell surface proteases, ADAM10 and urokinase (**Figure S4b**). Interesting, side-by-side comparison of proteins identified in CSP and secretomics did not show a correlation between neo-N-termini and protein level enrichment which would not necessarily be expected. Extracellular proteolysis is known to impact the protein turnover and stability of proteins displayed on the cell surface, which can blur this comparison (**Figure 4e**).

Lastly, we also found that hypoxia induces the release of a small number of proteins that typically reside within intracellular compartments such as the cytoplasm, lysosome, endoplasmic reticulum, or golgi apparatus (**Figure 4c**). It is possible these are the result of some cell lysis or other forms of cellular stress in hypoxia that leads to exocytosis of intracellular proteins. Further work will be needed to identify the mechanism underlying the release of these proteins into the media.

## 3. Discussion and Conclusion

Hypoxia is an important feature of the tumor microenvironment and greatly impacts the cellular and molecular fate of cells. PDAC tumors represent the extreme end of the hypoxic spectrum, where tumor cores can be nearly 25-times more hypoxic that normal tissues^4,10^ To identify neo-biomarkers and therapeutic opportunities for this disease, we sought to understand how hypoxia broadly affects the cell surface landscape. Leveraging mass spectrometry techniques suited for identifying extracellular modifications, we discovered that hypoxia strongly impacts proteolysis and protein shedding. Using CSP technology, we identified 906 N-termini on 384 membrane proteins that were either up or down-regulated in response to hypoxia. The vast majority corresponding to type I single-pass transmembrane proteins (**Figure 1b**); most cleavages occurred in the unstructured loops of protein ectodomains (**Figures 1c, 1d**). These results reflect a similar pattern we previously identified in breast cancer cells under normoxic conditions^22^, indicating that hypoxia can induce changes to the proteolytic landscape that are not dependent on protein levels. CSP revealed that PDAC cell surface protein abundance and proteolysis profiles vary under hypoxic conditions (**Figure S2a**). Like most cancers, PDAC is a complex disease with different subtypes^30,42^; the cell lines chosen for this study reflect this diversity. PaTu8902 cells express more epithelial differentiation genes, whereas Panc-1, MIA PaCa-2, and KP4 cells express more basal-like markers, such those involved in stem-cell and epithelial-to-mesenchymal transition (EMT)^42^. Additionally, these PDAC cell lines carry different KRAS mutations which may contribute to different activation levels of the Ras pathway and protease expression^43^. When peptides were narrowed down to those that have a >1.8-fold enrichment, either in normoxic or hypoxic conditions, we found that MIA-PaCa-2 cells displayed increased proteolysis at these sites in hypoxia, while the other three cell lines showed a general decrease in proteolysis for the same targets. These differences are reflected in the differences in proteases that are activated in each cell line. Further functional genomic or selective inhibitor studies may help to tease out which proteases are responsible for which specific proteolytic event.

There has been a surge of interest in developing conditionally active “masked” antibodies that exploits universal up-regulation of proteolysis in the TME. An antibody is designed to bind the protein of interest (POI) only when the binding site is exposed due to a proteolytic cleavage. This strategy provides a significant expansion of the therapeutic index as shown for masked anti-EGFR antibodies that spare healthy tissues that express the POI but lack active proteases, while attacking the cancer cells that over express the POI and upregulate proteolysis^60^. Our data help inform design principles for cleaving the protease mask. (i) Most natural cuts appear in loop regions to best imbed the cleavage site in a loop. (ii) the PDAC cancer cells studies here express and activate dozens of proteases so one can create a more generalized loop sequence and this is reinforced by the broad and degenerate sequence Logo we observe.

GO analysis revealed an over-representation of proteins involved in cellular adhesion and migration (**Figure 2c**). In particular, a change of >1.8 fold enrichment ratio (up or down) was seen for 45 different proteins involved in cell adhesion and signaling. Prominent among these were ITGB1^44,45^, ITGAV^45^, and ITGB5^45^, that are known to be upregulated in expression under hypoxic conditions and promote cellular migration. It is possible that downregulating the cleavage of these molecules may be integral to enhancing cell adhesion once the cells have metastasized. It has also been shown that cleavage of cell adhesion molecules can contribute to signal transduction^46,47^. The proteolytic ectodomain cleavage of CADH2 is involved in signal transduction and the degradation of CREB-binding protein (CBP), a transcriptional coactivator^47^. Exposure to hypoxia may trigger a different response in MIA PaCa-2 cells that results in increased proteolysis for signaling rather than adhesion. The western blot validation of RTN4R in particular demonstrates how hypoxia can push proteolysis in opposite directions in different cell lines (**Figure 2b**). We observed an increase in the cleavage product in PaTu 8902 cells while proteolysis was completely inhibited in the other three cell lines. Thus, it appears that proteolysis drives a complex circuit of events not easily explained by a single protease or cleavage event.

To complement our studies of the cell surface proteolytic landscape, we also investigated how hypoxia affected what proteins were shed or secreted into the media. 517 membrane proteins were identified in the culture supernatant and as with the N-terminomics data, type I single-pass transmembrane proteins were the largest class of identified proteins. As expected with a method tailored to isolating untethered glycoproteins or fragments, an increased portion of the identifications were secreted proteins (**Figure 3b**). Comparing the datasets across the four cell lines revealed three common clusters of proteins. In Cluster 1 (**Figure 3d**), GO analysis revealed an over-representation of proteins involved in regulating inflammation and the humoral immune response. Canonically, proteins in the complement system often defend against inflammation, however studies have shown that PDAC increases the expression of complement proteins that can then trigger pro-inflammatory cytokines^48,49^. In other contexts, hypoxia further increases expression of complement proteins^50,51^, but interestingly, we found that these proteins were downregulated, indicating that there are likely additional factors in the tumor microenvironment outside of hypoxia that influence complement expression in PDAC. Cluster 2 shows an upregulation of molecules that impact cellular metabolism. This corroborates existing data showing that hypoxia upregulates molecules such as TGF-β^52^ and alpha-enolase^53^ that are involved in hypoxia tolerance. Cluster 3 shows increased secretion of molecules involved in tissue development and an over-representation of protease inhibitors. Taking a closer look at the secreted proteases (**Figure 4a**) a moderate number that are overexpressed under hypoxic conditions are involved in degrading the extracellular matrix. Those involved in coagulation are generally downregulated. Interestingly, some of the proteases over expressed in cluster 3 are inhibitors to those found in **Figure 4a**. TFPI1^54^ and APLP2^55^ are inhibitors of coagulation factors X & XI (FA10, FA11); GDN^56^ inhibits thrombin and urokinase; and SPIT1^57^ inhibits matriptase (ST14). APLP2 is also known to be shed by ADAM10 and ADAM17^58^. Although a number of proteases are overexpressed and secreted into the supernatant, it is possible that the presence of the inhibitors is enough to reduce proteolysis seen in **Figure 3a**.

By comparing our CSP and secretomics data reflected membrane bound and secreted proteins, respectively, we identified 11 proteases identified by both approaches (**Figure S4**). However, only the activating cleavage sites for ADAM10 and urokinase have been identified by CSP analysis. While this is not an indication that the other proteases are inactive, this mass spec method can only confirm that ADAM10 and urokinase are active. We also compared the enriched N-terminomics dataset to their respective secreted protein levels (**Figure 4e**) and did not see significant correlation; hierarchical clustering maintained them as separate groups. This was supported by correlation plots of the data sets for each cell line (**Figure S5**). In cases where we observed shedding of the entire ectodomain, it is possible the remaining membrane tethered fragment may be rapidly degraded so not detected in the CSP data set.

In summary, our data show that hypoxia induces bi-directional changes in the proteolytic landscape that can be even larger than changes in expression levels for surface and secreted proteins. Our data reveal there is a complex network of protease activation (both membrane and secreted forms) and inhibitor expression. It is important to note that all the proteolytic events we found are endogenous to the cancer cell lines so likely to be cancer specific. Despite the complexity of individual proteolytic events, hypoxia induces similar GO terms related to cell adhesion and migration, which are both hallmarks of cancer. Neo-proteolytic events on POIs can be targeted with therapeutic antibodies and provide an additional “and” gate for selectivity and safety^59^. Our goal was to provide a global view of the proteolytic landscape on the cell surface and secretome. Clearly more biological validation of these proteolytic events is needed. Nonetheless, the identification of hypoxia induced neo N-termini could lead to the development of new targeted therapies with fewer off-target effects, and shed fragments may serve as new serum biomarkers in cancer.

---

**Supplementary Figure S1:**
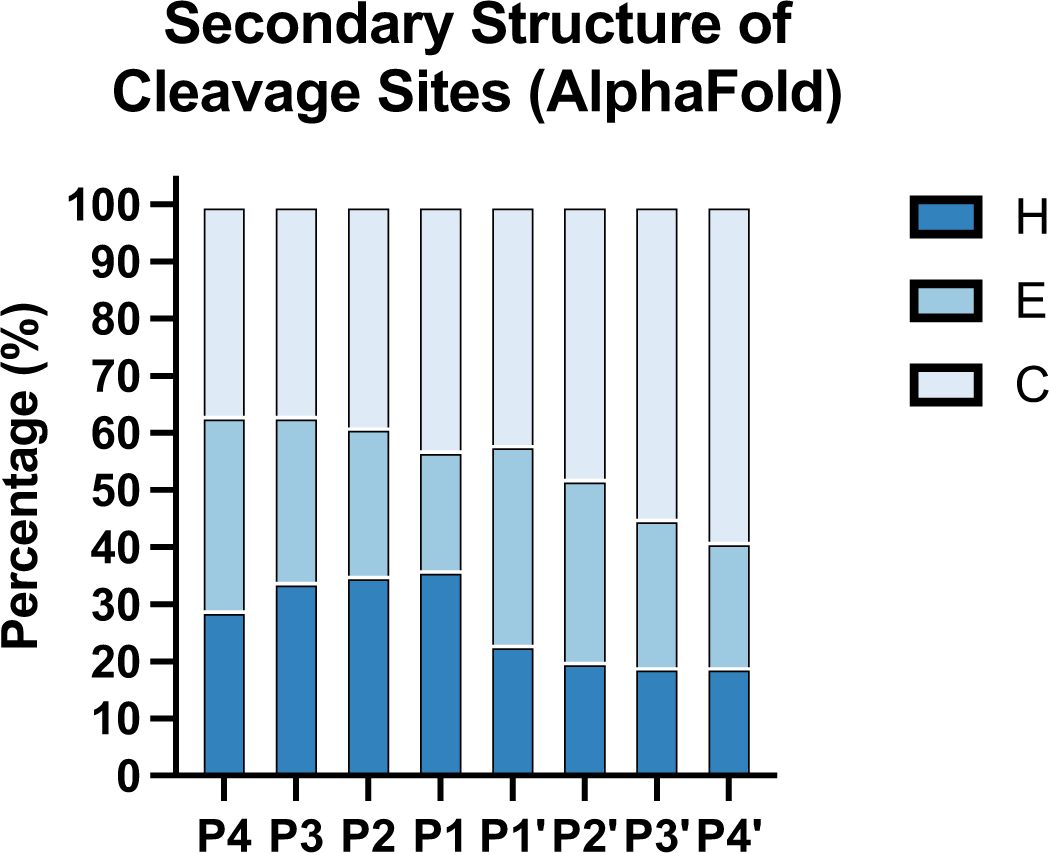
Predicted secondary structure at cleavage sites. For proteins that did not have a structure deposited on PDB, AlphaFold 2.0 models were used to predict the secondary structure surrounding the identified cleavage sites. H represents helices, E represents residues in sheets, and C represents residues in loop and unstructured locations.

**Supplementary Figure S2:**
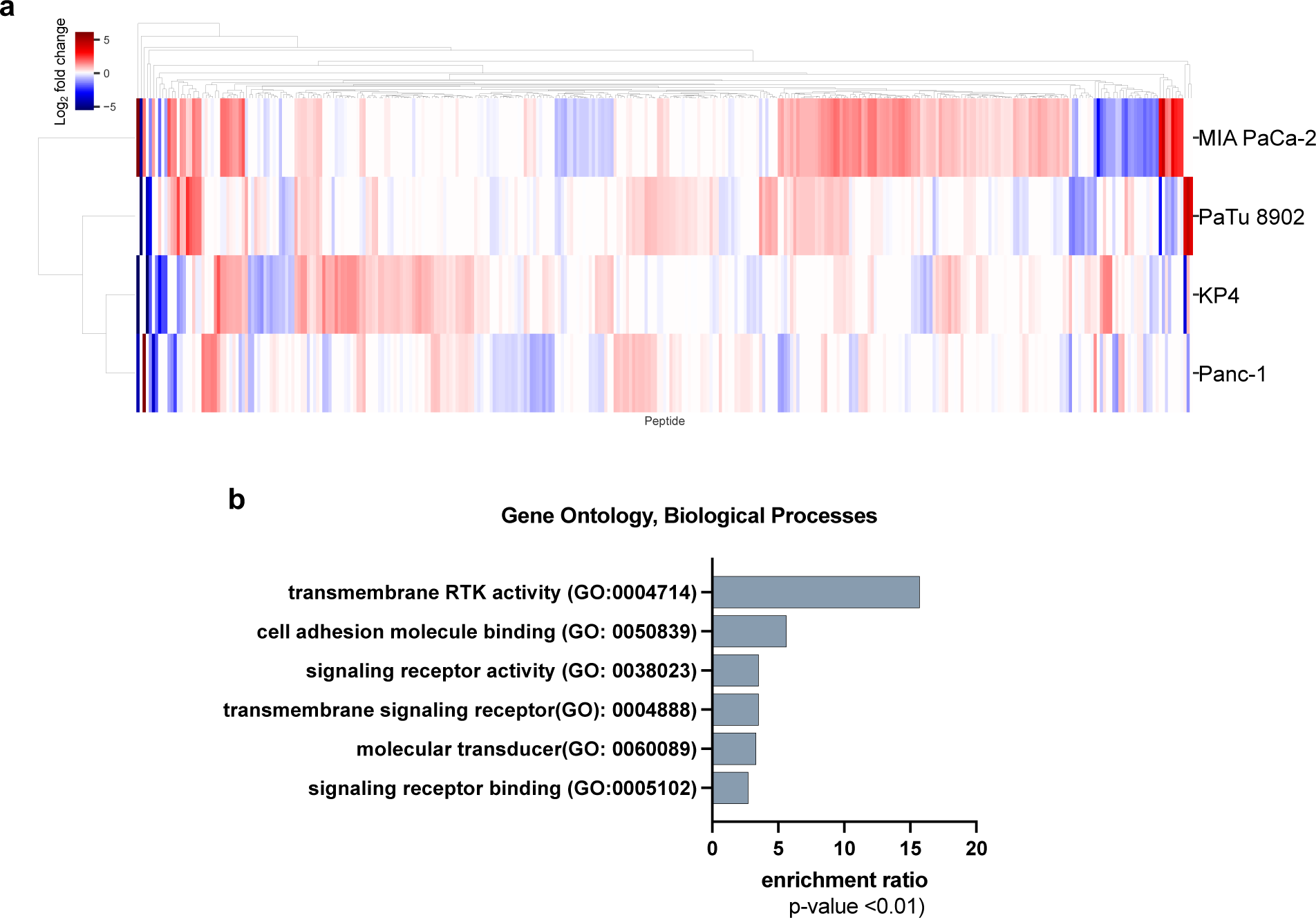
Quantification of N-termini abundance changes observed under normoxic conditions. **a.** Heatmap of identified N-termini depicting the change in peptide abundance under hypoxic conditions versus normoxic conditions. Red indicates an increased presence in hypoxia while blue indicates an increased presence in normoxia. **b.** GO analysis of all N-termini reveals an over-representation of proteins involved in signaling receptor activity.

**Supplementary Figure S3:**
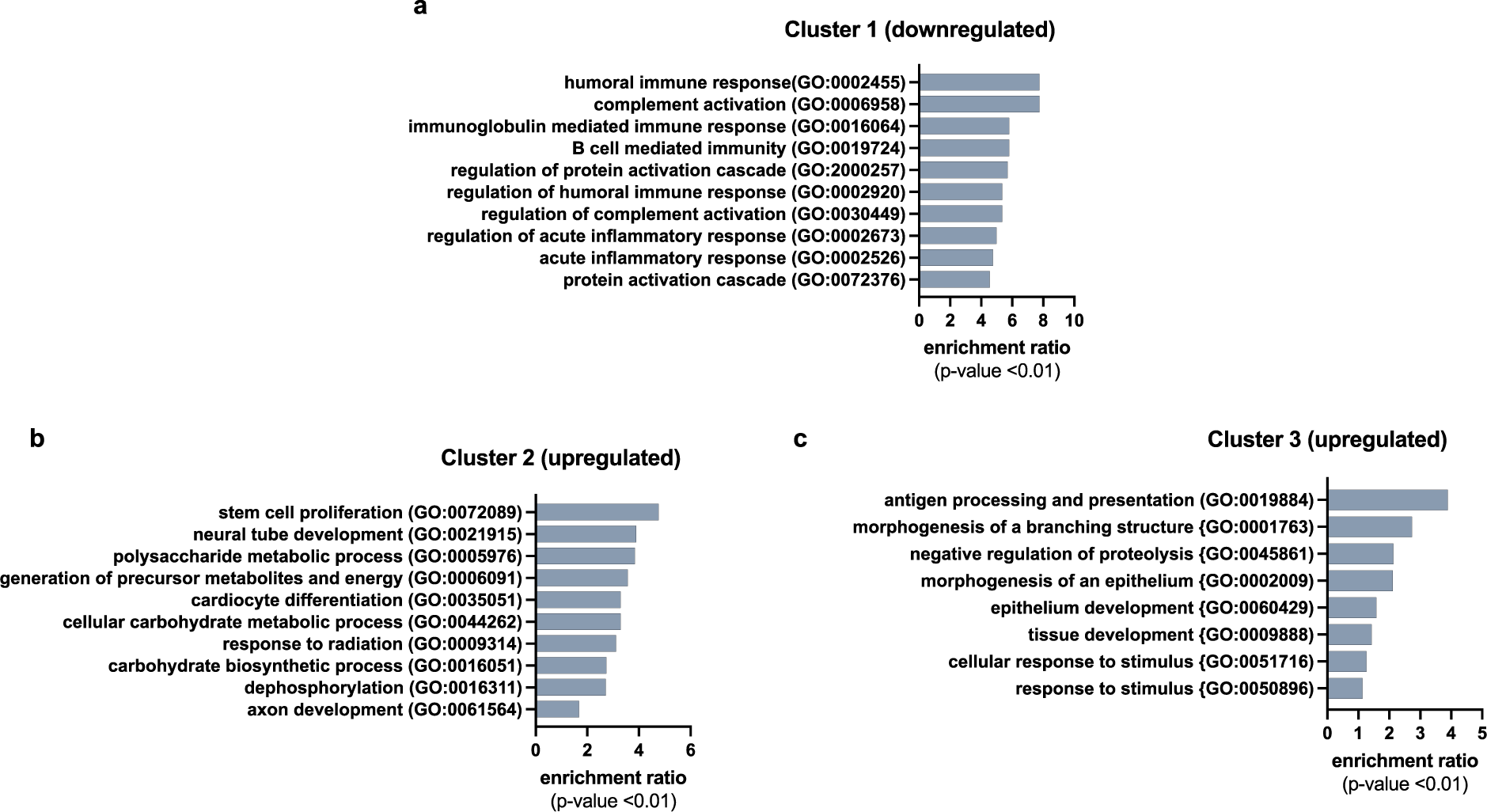
GO Analysis of secretomics. **a.** Cluster 1 identifies downregulated proteins involved in regulating the immune and inflammatory responses. **b.** Cluster 2 identifies moderately upregulated proteins involved in regulating cellular metabolism and development **c.** Cluster 3 identifies upregulated proteins involved in antigen processing/presentation, as well as those involved in tissue development and response to stimuli.

**Supplementary Figure S4:**
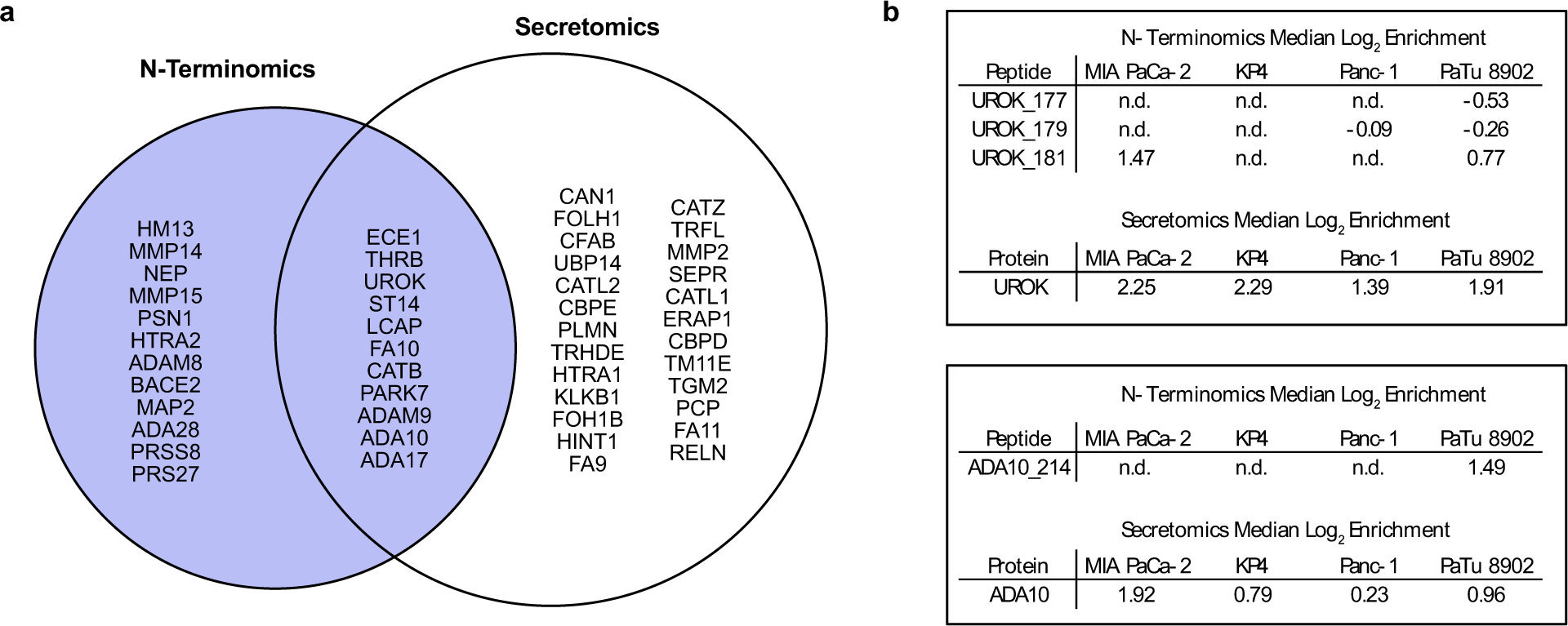
Shared proteases identified in N-terminomics and secretomics. **a.** Venn diagram displaying functionally-diverse proteases identified in N-terminomics, secretomics, and shared between MS approaches. **b.** Cleavages observed at activating sites of urokinase and ADA10 and their associated secreted protein levels with positive values reflecting more cut in hypoxia vs normoxia and negative values the opposite. n.d. = not detectable.

**Supplementary Figure S5:**
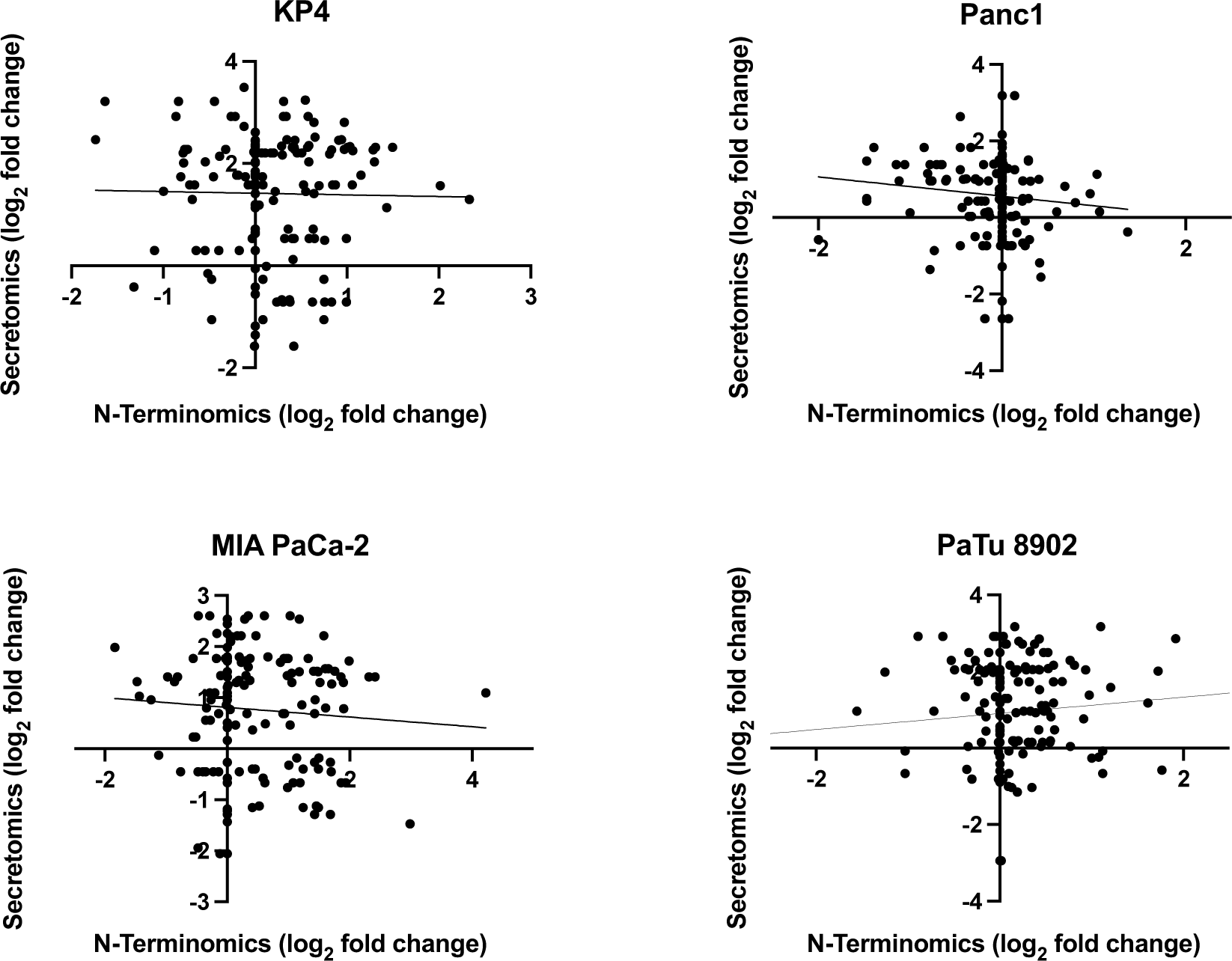
Correlation plots between N-terminomics and secretomics data. Identified N-termini (x-axis) and their changes under hypoxic conditions plotted against their respective change in protein levels in secretomics (y-axis) datasets show little correlation between proteolysis and shed/secreted proteins.

## Materials and Methods

### Cell Culture

PANC-1, and PaTu 8902 were purchased from ATCC (Manassas, Virginia), and MIA-PaCa and KP4 cell lines were gifted by the Perera lab at UCSF. Cell lines were tested for Mycoplasma contamination on a bi-yearly basis. All 4 cell lines were cultured in IMDM SILAC media (Thermo Fisher Scientific; Waltham, MA) + 10% dialyzed FBS (GeminiBio, Sacramento, CA) and 1% Penicillin/Streptomycin (Thermo Fisher Scientific) containing L-[^13^C_6_, ^15^N_2_] lysine and L-[^13^C_6_, ^15^N_4_] arginine for heavy labeling (Cambridge Isotype Laboratories, Tewksbury, MA) or L-{^12^C_6_, ^14^N_2_] lysine and L-[^12^C_6_, ^14^N_4_] arginine for light labeling (Sigma-Aldrich, St. Louis, MO). Under normoxic conditions, cells were grown at 37°C, 5% O_2_ for 72 hours before harvest. Under hypoxic conditions, cells were grown at 37°C, 1% O_2_ for 72 hours before harvest.

### Cell Surface Capture Proteomics

Labeling and capture of cell surface proteins has been described in detail previously (Ref. 61). Cells were harvested with Versene (Thermo Fisher Scientific). Cells were washed with phosphate buffered saline pH6.5 (Cytiva, Marlborough, MA) and incubated with WGA peroxidase (Vector Labs, Newark, CA) and biotin tyramide (Apex Bio, Houston, TX) at 37°C. Hydrogen peroxide (Sigma-Aldrich) was added and the reaction allowed to proceed for 2 minutes before being quenched and washed with sodium pyruvate (Cytiva). Cells were collected and lysed in RIPA buffer with protease inhibitor cocktail (Sigma-Alrich). Cell lysate was incubated with neutravidin agarose beads (Thermo Fisher Scientific at 4°C for 30 minutes. Beads were washed with RIPA buffer, 50 mM PBS + 1M NaCl, and 50 mM ammonium bicarbonate +2M urea buffer. The washed beads were processed for tryptic peptide elution and mass spectrometry analysis using the iST desalting kit (Preomics, Martinsried, Germany) per manufacturer instructions.

### Secretomics Capture

Secretome analysis has been described in detail previously [REF 23]. Cells were incubated under normoxic or hypoxic conditions with 100 μM N-Azidoacetymannosamine (Thermo Fisher Scientific) for 72 hours. Conditioned media was collected incubated with Concanavalin A Agarose beads (G-Biosciences, Overland, MO) for 2 hours at room temperature. Proteins were eluted from the ConA beads with methyl-alpha-mannopyranoside (Sigma-Aldrich) and incubated with DBCO beads (Vector Laboratories; Newark, CA) overnight at 4°C. Protein disulfide bonds were reduced with DTT (GoldBio, Olivette, MO) and cysteines alkylated with iodoacetamide (Sigma-Alrich). The protein was digested with sequencing grade trypsin and the tryptic peptides were processed for mass spectrometry analysis using the iST desalting kit per manufacturer instructions.

### Cell Surface N-terminomics

N-terminomics was performed as previously described [ref 22]. Cells were harvested with Versene and treated with sodium periodate (Sigma-Aldrich) to oxidize cell surface glycans. Aminooxy-peg2-stabiligase and analine (Sigma-Alrdich) were then added to cells to tether stabiligase to the cell surface for 15 minutes at 4°C. After washing, cells were incubated at room temperature with a biotinylated peptide ester substrate for 15 minutes and washed with PBS. Cells were collected and lysed in RIPA buffer with protease inhibitor cocktail. Cell lysate was incubated with neutravidin agarose beads at 4°C for 30 minutes. Beads were then washed with RIPA buffer, 50 mM PBS + 1M NaCL, and 50 mM ammonium bicarbonate +2M urea buffer. The beads were then incubated with sequencing grade trypsin (Promega, Madison, WI) overnight at room temperature to release tryptic peptides. Beads were collected and washed with 50 mM ammonium bicarbonate buffer and incubated with TEV protease and DTT overnight. The TEV protease elution was collected and processed for mass spectrometry analysis using the iST desalting columns per manufacturer instructions.

### Liquid chromatography mass spectrometry analysis

200 ng of prepared samples were injected onto a 25 cm, ReproSil c18 1.5 uM 100 A column (PepSep) on a timsTOF Pro with a Captive Spray source and a nenoElute line (Bruker; Hamburg, Germay). a stepwise linear gradient method with H2O in 0.1% Formic acid and acetronitrile with 0.1% formic acid (solvent B): 5-30% solvent B for 90 min at 0.5 µl/min, 30-35% solvent B for 10 min at 0.6 µl/min, 35-95% solvent B for 4 min at 0.5 µl/min, 95% hold for 4 min at 0.5 µl/min) was used. Acquired data was collected in a data-dependent acquisition (DDA) mode with ion mobility activated in PASEF mode. Data was then analyzed using PEAKS online Xpro 1.6 (bioinformatics Solutions Inc.; Ontario, Canada), using the SwissProt GOCC Plasma Membrane Database. Analyzed data was further processed using a python script previously described^22^. Raw data files have been deposited to the ProteomeXchange consortium with the dataset identifier PXD057350.

### Western Blots

20 μg of cellular lysates were loaded onto NuPAGE 4%-12%, Bis Tris Mini Protein Gels (Thermo Fisher Scientific). Gels were then transferred to PVDF membranes using the iBlot2 (Thermo Fisher Scientific). Membranes were blocked with 5% bovine serum albumin (BSA; Gemini Bio) for 1 hour at room temperature. Primary antibodies were added to the blocking buffer and incubate at 4°C overnight. β-actin (1:2000), CDCP1 (1:1000), and MUC-18 (1:500) antibodies were purchased from Cell Signalling Technology (Danver, MA). The RTN4R (1:250) antibody was purchased from Proteintech (Rosemont, IL). Membrane was then washed 3X with PBST and incubated with LI-COR secondary antibodies (Lincoln, NE) for 1 hour at room temperature. Membrane was washed three times and imaged on the Odyssey DLx system (LI-COR).

## REFERENCES

1. Siegel, R.L., Gianquinto, A.N., & Jemal, A. Cancer Statistics, 2024. CA Cancer J. Clin 74, 12–49 (2024).

2. Murakami, T., Hiroshima, Y., Matsuyama, R., Homma, Y., Hoffman, R., & Endo, I. Ann. Gast. Surg. 2, 130–137 (2019).

3. Sherman, M.H. & Beatty, G.L. Annu. Rev. Pathol. Mech. Dis. 18,, 123–148 (2022).

4. Muz, B., de la Puente, P., Azab, F., & Azab, A.K. Hypoxia 3, 83–92 (2015).

5. Li, Y., Zhao, L., Li, & X-F. Technol. Cancer Res. Treat. 20 (2021).

6. Chen, Z., Han, F.m Du, Y., Shi, H., & Zhou, W. Sig. Transduct. Target Ther. 8 (2023). a. Ruan, K. J. Cellular Biochemistry (2009)

7. Wang, B., Zhao, Q., Zhang, Y., Liu, Z., Zheng, Z., Liu, S., Meng, L., Xin, Y. & Jiang, X. J. Exp. Clin. Canc. Res. 40 (2021).

8. Damgaci, S., Ibrahim-Hashim, A., Enriquez-Navas, P.M., Pilon-Thomas, S., Guvenis, A., & Gillies, R.J. Immunology 154, 354–362 (2018).

9. Khouzam, R.A., Goutham, H.V., Zaarour, R.F., Chamseddine, A.N., Francis, A., Buart, S., Terry, S., & Chouaib, S. Semin. Cancer Biol. 65, 140–154 (2020).

10. Koong, A.C., Mehta, V.K., Le, Q.T., Fisher, G.A., Terris, D.J., Brown, J.M., Bastidas, A.J., & Vierra, M. Int J. Radiat Oncol. Biol. Phys. 48, 919–922 (2000).

11. Vizovisek, M., Ristanovic, D., Menghini, S., Christiansen, M.G., & Schuerle, S. Int J. Mol. Sci. 22, 2514–2533 (2021).

12. Turk, B., Turk, D, & Turk, V. EMBO J. 4, 1630–1643 (2012).

13. Werb, Z. Cell 91, 439–442 (1997).

14. Wolf, D.H. & Menssen, R. FEBS Letters 592, 2515–2524 (2018).

15. Dudani, J.S., Warren, A.D., & Bhatia, S.N. Annu. Rev. Canc. Biol. 2, 353–376 (2018).

16. Eatemadi, A., Aiyelabegan, H.T., Negahdari, B., Mazlomi, M.A., Daraee, H., Daraee, N., Eatemadi, R., & Sadroddiny, E. Biomed. Pharmacother. 86, 221–231 (2017).

17. Griswold, A.R., Cifani, P., Rao, S.S., Axelrod, A.J., Miele, M.M., Hendrickson, R.C., Kentsis, A., & Bachovchin, D.A. Cell Chem. Biol. 26, 907–907 (2019).

18. Staes A., Damme, P., Timmerman, E., Ruttens, B., Stes, E., Gevaert, K., & Impens, F. Methods Mol Biol 1574, 51–76 (2017).

19. Prudova, A., Gocheva, B., Auf dem Keller, U., Eckhard, U., Olson, O.C., Akkari, L., Butler, G.S., Fortelny, N., Lange, P.F., Mark, J.C., Joyce, J.A., & Overall, C.M. Cell Rep. 16, 1762–1773 (2016).

20. Dix, M.M., Simon, G.M., & Cravatt, B.F. Methods Mol Biol. 1113, 6–70 (2014).

21. Weeks, A.M. & Wells, J.A. Curr. Protoc. Chem. Biol. 12 (2020).

22. Schaefer, K.S., Lui, I., Byrnes, J.R., Kang, E., Zhou, J., Weeks, A.M., & Wells, J.A. ACS Centr. Sci. 8, 1447–1456 (2022).

23. Tüshaus, J., Müller, S.A., Kataka, E.S., Zaucha, J., Monasor, L.S., Su, M., Güner, G., Jocher, G., Tahirovic, S., Frishman D., Simons, M., & Lichtenthaler, S.F. EMBO J. 15 (2020).

24. Meissner, F., Scheltema, R.A., Mollenkopf, H-J., & Mann, M. Science 340, 475–478 (2013).

25. Dieterich, D.D., Link, A.J., Graumann, J., Tirrell, D.A., & Schuman, E.M. PNAS 103, 9482–9487 (2006).

26. Eichelbaum, K., Winter, M., Berriel Diaz, M., Herzig, S., & Krijgsveld, J. Nat. Biotechnol. 30, 984–990 (2012).

27. Sun, J.S., Zhang, X.L., Yang, Y.J., & Nie, Z.G. Oncol. Lett. 10, 835–840 (2015).

28. Jiang, J., Tang, Y.L. & Liang, X.H. Canc. Biol. Ther. 11, 714–723 (2011).

29. Chen, S., Chen, J.Z., Zhang, J.Q., Chen, H.X., Yan, M.L., Huang, L., Tian, Y.F., Chen, Y.L., & Wang, Y.D. Canc. Lett. 383, 73–84 (2016).

30. Collison, E.A., Sadanandam, A., Olson, P., Gibb, W.J., Truitt, M., Gu, S., Cooc, J., Weinkle, J., Kim, G.E., Jakkula, L., Feiler, H.S., Ko, A.H, Olshen, A.B., Danenberg, K.L., Tempero, M.A., Spellman, P.T., Hanahan, D., & Gray, J.W. Nat. Med. 17, 500–503 (2011).

31. Werba, G., Weissinger, D., Kawaler, EA., Zhao, E., Kalfakakou, D., Dhara, S., Wang, L., Lim, H.B., Oh, G., Jing, X., Beri, N.m Khanna, L., Gonda, T., Pberstein, P., Hajdu, C., Loomis, C., Heguy, A., Sherman, M.H., Lund, A.W., Welling, T.H., Dolgalev, I., Tsirigos, A., & Simone, D.M. Nat. Commun. 14 (2023).

32. Swietlik, J.J., Bärthel, S., Falcomatà, C., Fink, D., Sinha, A., Cheng, J, Ebner, S., Landgraf, P., Dieterich, D.C., Daub, H., Saur, D., & Meissner, F. Nat. Commun. 14 (2023).

33. Lash, G.E., Fitzpatrick, T.E., & Graham, C.H. Biochem. Biophys. Res. Commun. 287, 622–629 (2001).

34. Crossin, K.L. Cell Adh. Migr. 6, 49–58 (2012).

35. Fei, M., Guan, J., Xue, T., Qin, L., Tabg, C., Cui, G., Wang, Y., Gong, H., & Feng, W. Cell. Mol. Biol. Lett. 23 (2018).

36. Saxena, K., Jolly, M.K., & Balamurugan, K. Transl. Oncol. 13 (2020).

37. Semenza, G.L. Physiology 36, 73–83 (2021).

38. Khouzam, R.A., Zaarour, R.F., Brodaczewska, K., Azakir, B., Venkatesh G.H., Thiery, J, Terry, S., & Chouaib, S. Front. Immunol. 13 (2022).

39. Eales, K.L., Hollinshead, K.E.R., & Tennant D.A. Oncogenesis 5, e190 (2016).

40. Kierans, S.J. & Taylor, C.T. J. Physiol. 599, 23–37 (2021).

41. Frezza, C., Zheng, L., Tennant, D.A., Papkovsky, D.B., Hedley, B.A., Kalna, G., Watson, D.G., & Gottlieb, E. PLos ONE 6, e24411 (2011).

42. Adams, C.R., Htwe, H.H., Mash, T., Wang, A.L., Montoya, M.L., Subbaraj, L., Tward, A.D., Bardeesy, N., & Perera, R.M. eLife 8, e45313 (2019).

43. Johnson, C., Burkhart, D.L., & Halgis, K.M. Canc. Discov. 12, 913–923 (2022).

44. Ju, J.A., Godet, I., Ye, I.C., Byu, J., Jayatilaka, H., Lee, S.J., Xiang, L., Samanta, D., Lee, M.H., Wu, P., Wirtz, D., Semenza, G.L., & Gilkes, D.M. Mol. Cancer. Res. 15, 723–734 (2017).

45. Befani, C. & Liakos, P. Cell Biol. Intl. 41, 769–778 (2017).

46. Nagappan-Chettiar, S., Johnson-Venkatesh, E.M., & Umemori, H. Neurosci Res. 116, 60–69 (2016).

47. Reiss, K., Maretzky, T., Ludwig, A., Tousseyn, T., de Strooper, B., Hartmann, D., & Saftig, P. EMBO J. 24, 742–752 (2005).

48. Afshar-Kharghan, V. J. Clinic. Invest. 127, 780–789 (2017).

49. Hussain, N., Das, D., Pramanik, A., Pandey, M.K., Joshi, V., & Pramanik, K.C. Cancer Drug Resist. 5, 317–327 (2022).

50. Mueller-Buehl, A.M., Buehner, T., Pfarrer, C., Deppe, L., Peters, L., Dick, B.H., & Joachim, S.C. Cells 10, 3575 (2021).

51. Khan, M.A., Shamma, T., Kazmi, S., Altuhami, A., Ahmed, H.A., Assiri, A.M., & Broering, D.C. J. Transl. Med. 18, 147 (2020).

52. Mallikarjuna, P., Zhou, Y., & Landström, M. Biomolecules 12, 635 (2022).

53. Sedoris, K.C., Thomas, S.D., & Miller, D.M. BMC Cancer 10, 157 (2010).

54. Mast, A.E. & Ruf, W. J. Thromb Haemost. 20, 1290–1300 (2022).

55. Xu, F., Previti, M.L., Nieman, M.T., Davis, J, Schmaier, A.H., & Van Nostrand, W.E. J. Neurosci. 29, 5666–5670 (2009).

56. Festoff, B.W., Rap, J.S., & Chen, M. Neurology 42, 1361 (1992).

57. Skovbjerg, S., Holt-Danborg, L., Nonboe, A.W., Hong, Z., Frost, A.K., Schar, C.R., Thomas, C.C, Vitved, L., Jensen, J.K., & Vogel, L.K. Biochem J. 477, 1779–1794 (2020).

58. Endres, K., Postina, R., Schroeder, A., Mueller, U., & Fahrenholz, F. FEBS J. 272, 5808–5820 (2005).

59. Lim S.A., Zhou J., Martinko A.J., Wang Y.H., Filippova E.V., Steri V., Wang D., Remesh S.G., Liu J., Hann B., Kossiakoff A.A., Evans M.J., Leung K.K., Wells J.A. J Clin Invest. 132(4):e154604 (2022)

60. Desnoyers L.R., Vasiljeva O.., Richardson J.H., Yang A., Menendez E.E., Liang T.W., Wong C., Bessette P.H., Kamath K., Moore S.J., Sagert J.G., Hostetter D … R, Han F, Gee J., Flandez J., Markham K., Nguyen M., Krimm M., Wong K.R., Liu S., Daugherty P.S., West J.W., Lowman H.B. Sci Transl Med. 5(207):207ra144 (2013)

61. Kirkemo L.L., Elledge S.K., Yang J., Byrnes J.R., Glasgow J.E., Blelloch R., Wells J.A. Elife 11:e73982 (2022)

